# Truvari: Refined Structural Variant Comparison Preserves Allelic Diversity

**DOI:** 10.1101/2022.02.21.481353

**Authors:** Adam C. English, Vipin K. Menon, Richard Gibbs, Ginger A. Metcalf, Fritz J. Sedlazeck

**Affiliations:** Baylor College of Medicine Human Genome Sequencing Center, Houston, TX, USA

**Keywords:** Structural Variation, SV Comparison, SV Merging, SV Benchmarking, SV Annotation

## Abstract

For multi-sample structural variant analyses like merging, benchmarking, and annotation, the fundamental operation is to identify when two SVs are the same. Commonly applied approaches for comparing SVs were developed alongside technologies which produce ill-defined boundaries. As SV detection becomes more exact, algorithms to preserve this refined signal are needed. Here we present Truvari - a SV comparison, annotation and analysis toolkit - and demonstrate the effect of SV comparison choices by building population-level VCFs from 36 haplotype-resolved long-read assemblies. We observe over-merging from other SV merging approaches which causes up to a 2.2x inflation of allele frequency relative to Truvari.

## Background

The march of progress in genomic sequencing is constant, with increasing speed from improving technologies being applied to growing populations/cohorts and leading to discoveries from increasingly harder to assess genomic regions. One particular example of this progress over the last two decades has been the analysis of Structural Variants (SVs) which includes 50bp or larger genomic alterations of the genome. While single-nucleotide variants vastly outnumber SVs by count, the cumulative number of bases altered by SVs is higher due to their size, resulting in a documented impact on disease development and progression[1–3].

The detection of SVs has been enhanced most notably through the advent of long-read sequencing. No longer hindered by alignment through the repetitive elements which frequently mediate SVs[4], long-read sequencing has enabled refined characterization of SVs[5]. Simultaneously, SV benchmarking standards such as that created by the Genome in a Bottle consortium (GIAB) have provided objective measurements on the quality of SV tools which has assisted both genome researchers and software developers[6]. However, these improvements have largely focused on SV discovery and genotyping within the context of a single sample[7]. When comparing SVs across multiple samples, the question of how best to identify matching SVs remains inadequately addressed.

SV comparison is a fundamental operation of benchmarking, annotation, and merging that’s required to address both technical artifacts and biological differentiation. First, when SVs are called by different sequencing experiments or heterogeneous pipelines, any combination of base-calling errors[7], differences in pipeline sensitivity[5], and alignment ambiguities around repeats[8] may cause the same SV to be placed in different positions or contain different sequences. Furthermore, SV comparison choices impact evaluations of results. If SV matching is too lenient, benchmarking performance is inflated, incorrect annotations are applied, or over-merging occurs and causes unique SVs to be falsely identified as shared between samples. Over-merging is particularly problematic for merging due to losing allelic diversity and over-estimating allele frequency. Similarly, if SV comparison is too strict and matching SVs aren’t identified, benchmarking performance is deflated, annotations are missed, and experiments such as association analyses may become under-powered[9].

Multiple strategies for SV comparison have been proposed. For example, SVs are considered to be equal using reciprocal-overlap if a proportion of their individual sizes are overlapping. This traditionally has been applied to CNV calling (e.g. array-CGH) as the breakpoints are imprecise[10]. However, reciprocal-overlap is not applicable to sequence-resolved insertions, which have no physical span over the reference. With more precise breakpoints, other heuristics have been postulated such as breakpoint agreement where SVs are considered matching when their breakpoints are within a certain interval (e.g. 500-1000bp). While this method may generally be sufficient for larger SVs, it is insensitive to subtle differences of smaller SVs or those at complex loci with multiple events. The logical progression is to also take into consideration the length of the SV to improve the threshold/wobble distance allowance for the breakpoints[11–13]. However, insertions of the same length and at the same position may vary in sequence composition. Any of these approaches, in isolation, can incorrectly identify alleles as matching.

These concerns expose the need for a systematic approach to SV comparison that starts with a high-quality set of SV calls and builds from that an understanding of the impact of SV comparison choices. To accomplish this, we built Truvari, which assists SV comparison by leveraging multiple metrics to make informed comparisons choices. We use the experiences gained from Truvari being a widely used and recommended benchmark tool for SVs[6]. Truvari’s comparison approach is especially relevant given the improvements on SV calling accuracy in terms of breakpoint-exact, sequence resolved calls that are becoming commonplace, not only from long-read sequencing but also more exact short-read SV discovery algorithms[14,15].

We take previously published data of haplotype-resolved assemblies from 36 diverse individuals and measure the intra-sample haplotype similarity of SVs using Truvari[16,17]. We demonstrate how even high quality pipelines can produce similar, but not identical SV representations. These results are important to understand the impact of different methodologies on population merging. We again leverage Truvari to build pVCFs and gain insights into how SV merging choices affect biologically relevant metrics such as SV count and allele-frequency. We give these insights context by using Truvari with varying matching thresholds of these metrics as well as comparison to other SV merging methodologies.

## Results

### Truvari description

Truvari is an open source toolkit for the comparison, annotation, and analysis of Structural Variation. This research focuses on the SV comparison tools for benchmarking (*bench*) and merging (*collapse*) but leverages the annotation and analysis features to enrich the information presented. Truvari’s comparison approach is detailed in Methods (**Figure 1**). Briefly, Truvari compares SVs inside variant call format files (VCF) by measuring five similarity metrics between all pairs of calls within a region. These metrics are SVTYPE matching, reference distance, reciprocal-overlap, size similarity, sequence similarity and genotype matching. If any of the metrics violate user-defined thresholds, the pair of calls fails to be a candidate match.

**Figure 1:**
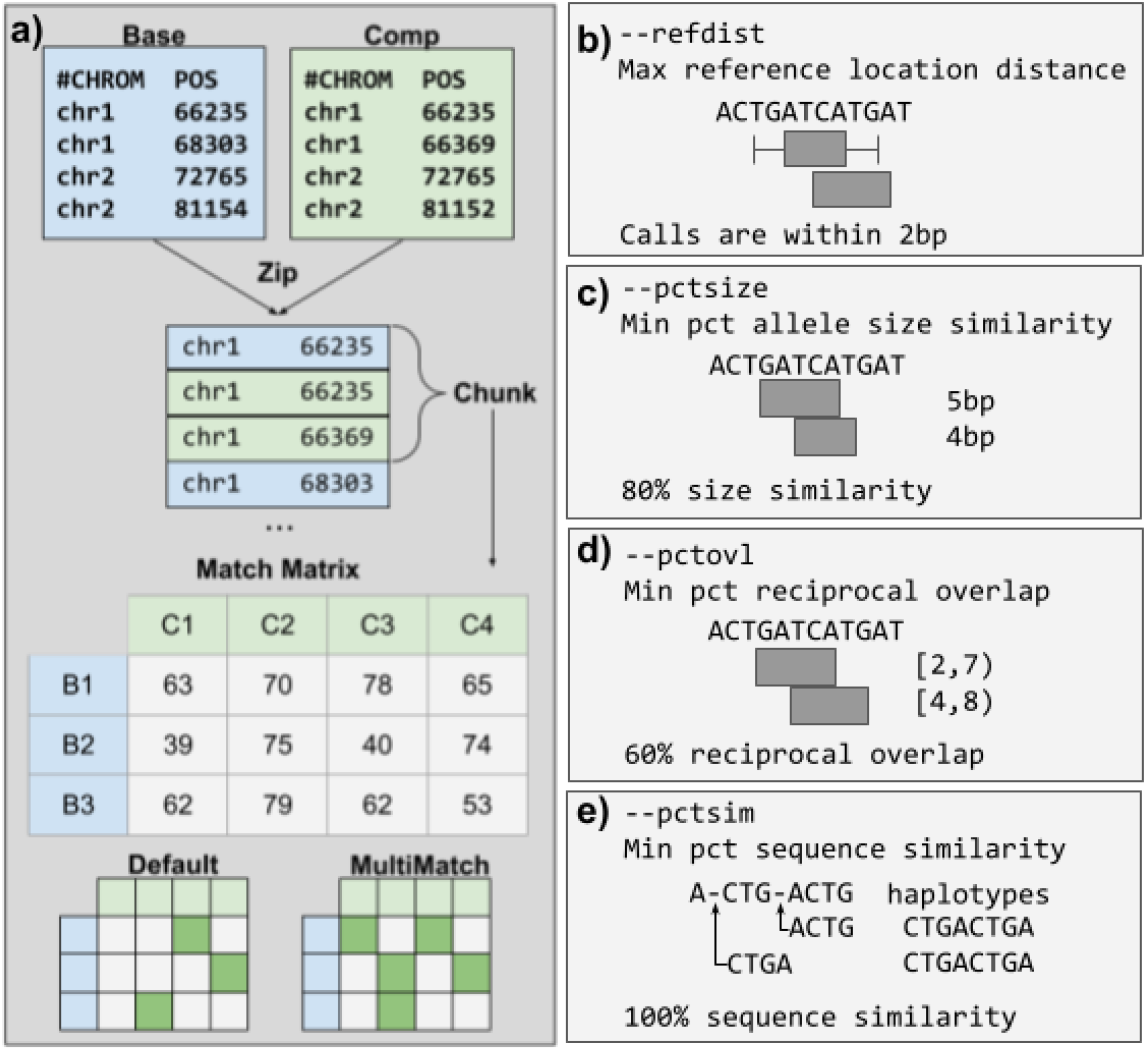
Overview of Truvari method and comparison metrics. a) Schematic illustrating the Truvari bench matching approach of a base and comparison (comp) VCF. b-e) Comparison metrics used by Truvari to measure similarity.

For Truvari *bench*, default matching thresholds are set to 70% sequence and size similarity, 500bp reference distance, svtype matching, and 0% reciprocal overlap. These thresholds are generally applicable to most single sample comparisons of a replicate to a ground-truth set of SVs. However, the thresholds can be raised or lowered based on the resolution of the SVs and desired stringency. For example, sequence similarity can be set to zero in order to capture matches between non-sequence-resolved calls. Truvari *collapse* default thresholds are 95% sequence and size similarity, 500bp reference distance, identical svtype, and 0% reciprocal overlap. These default thresholds work for highly-similar sequence resolved calls (e.g. calls from a harmonized pipeline) across multiple samples, but again can be tweaked to a user’s specifications.

### Matching SVs between haplotypes

To approach the central question of when to match a pair of SVs, we start with a set of 36 previously established, haplotype-resolved, long-read assemblies and call SVs[16,17]. We called SVs against three references - hg19, GRCh38 and the newly published chm13 - to observe how references impact the calling and analysis of the SVs[18–20]. First, to ensure a high accuracy of SV calling, we compared the NA24385 sample on hg19 against Genome in a Bottle (GIAB) v0.6 Tier1 SVs using Truvari *bench* (see methods). This measured a high precision (0.93) across each of the two haplotypes. Over 90.2% of true-positive SVs have at least 95% sequence similarity and size similarity between the generated calls and the GIAB truth set. This indicates highly consistent SV representations and that the SV calling methodology generated an accurate initial call set.

The simplest case of SV merging would be to combine SVs across haplotypes within a sample to create a diploid call-set. At most we expect a single match between haplotypes at homozygous alleles. For NA24385 on hg19, 5,478 SVs from each haplotype have identical sequence and position and therefore comprise homozygous alleles. The remaining 20,719 SVs were compared using Truvari *bench* to identify the similarity of SVs between haplotypes (**Figure 2a**). This showed 1,576 SV pairs having at least 95% sequence and size similarity and 1,195 between 70% to 95% similarity, all of which are candidates for merging. Interestingly, 402 SV pairs have ≤5% reciprocal-overlap but ≥70% sequence and size similarity. These pairs may indicate alignment ambiguities across repetitive regions (e.g. left-shift vs. right-shift).

**Figure 2:**
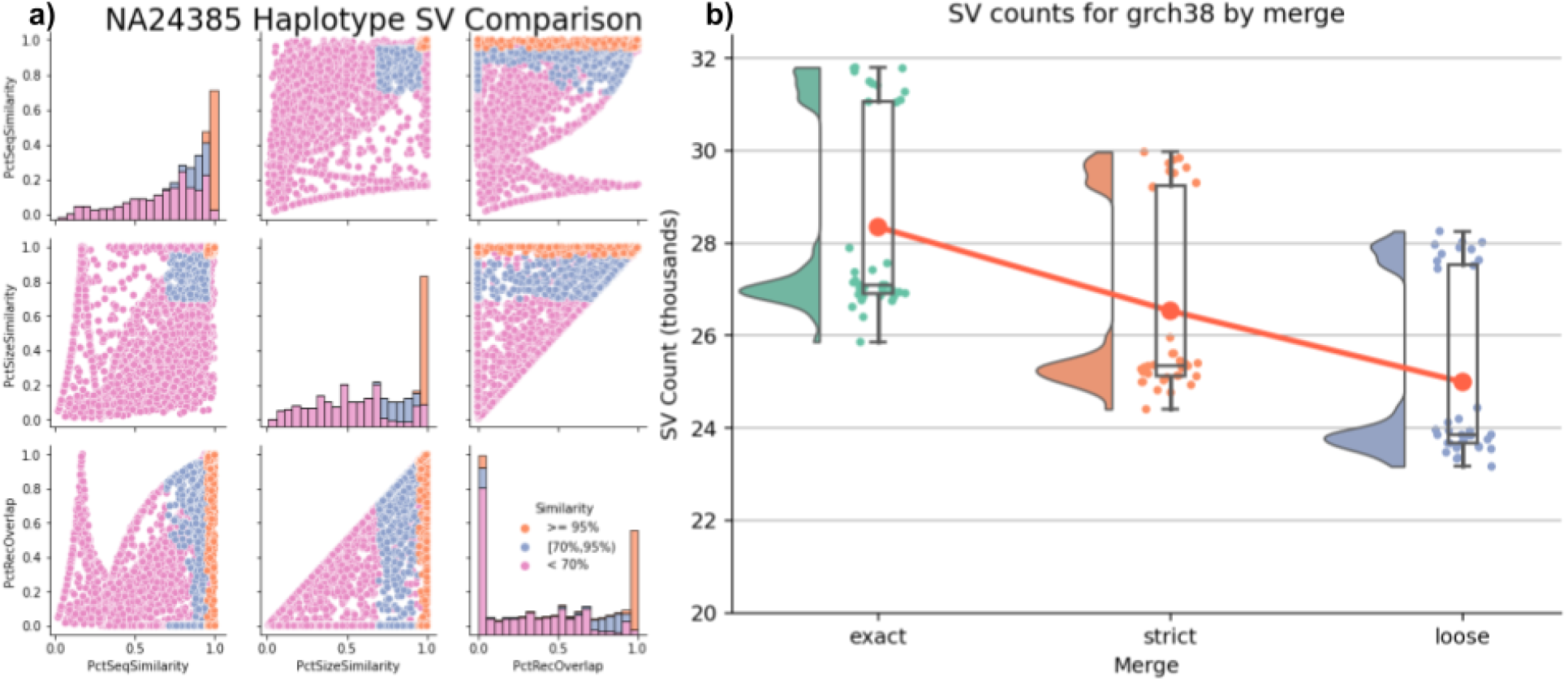
Intra-sample merging. a) Distributions of similarity metrics of SVs between NA24385 haplotypes. Colors are thresholds for sequence and size similarity. b) Effect of stringency on intra-sample merging SV counts for GRCh38, Trend line is the average number of SVs per-merge. Separation of samples is attributable to ancestry.

We investigated the effects of matching stringency on SV merging by creating three different merges: i) Exact method: the most stringent approach, combines SV if their breakpoint, size and sequence are identical. ii) Strict method: variants within 500bp and over 95% sequence and size similarity are merged; iii) Loose method: variants within 1,000bp and over 70% similarity are merged. We used Truvari *bench* to compare the three NA24385 intra-sample merges to GIAB Tier1 SVs (Supplementary Table 1). We observed 93.7% recall for the Exact and Strict merges, and 93.6% recall for Loose. The ratio of how many true-positive (TP) GIAB SVs are lost and how many potentially redundant calls are removed compared to the Exact merge is 1:790 for Strict and 8:1127 for Loose. Only 4.1% of false-negatives (FN) and 2.8% of false-positives (FP) from Strict have no complementary calls within 1,000bp. Therefore, with extremely permissive thresholds, Strict could have up to ∼96% recall and ∼97% precision. The remaining 391 FNs (238 INS; 153 DEL) are partially explained by 37% lacking aligned coverage from the assemblies as well as being enriched for SVs >=5,000bp (chi-square P<1E-5).

As merging becomes more permissive, SVs are more likely to find a match between haplotypes, thus lowering the overall SV count (**Figure 2b**). When looking across all 36 samples and all references, Exact produces an average of 27,187 SVs per-sample. Whereas the Strict and Loose merging lowers the average SV count by 1,520 and 2,851, respectively. Additionally, merging impacts the het/hom ratio due to heterozygous calls collapsing into a single homozygous variant (Supplementary Figure 1). The average and standard-deviation of het/hom ratios across samples and references are 4.9±0.9 for Exact merges, 3.2±0.7 for Strict, and 2.3±0.5 for Loose.

These patterns of merging’s effects on QC metrics appear on each reference and by SV types, though to differing degrees. For GRCh38 we see more SVs per-sample (26.6K) than chm13 (24.4K). However, GRCh38 has an imbalance of SV type frequency, with more insertions (16.5K) than deletions (10.1K), whereas chm13 is almost balanced (11.9K DEL, 12.4K INS). The most drastic change in SV counts due to merging comes from GRCh38 insertions where Loose merging results in a 15.9% decrease in SV count compared to Exact (Supplementary Figure 2). As previously reported[21], we observe a greater number of SVs from individuals of African ancestry with an average of ∼30.8K SVs compared to ∼25.6k SVs from all other individuals (Supplementary Figure 3).

This analysis of haplotype merging indicates the need for SV merging which attempts to remove redundant variant representations. The 95% sequence and size similarity thresholds from Strict merge has a well-balanced preservation of unique SVs and reduction of redundant SVs across individual samples for this call-set. Thus we chose these thresholds for Truvari *collapse* to produce the final per-sample VCFs.

### Inter-sample SV merging

Merging within a sample is relatively simple due to the pairing of haplotypes. However, merging between samples requires evaluating clusters of SVs. Next, we investigated how merging approaches perform across multiple samples. From the individual VCFs produced in the previous step, we created a project-level VCF (pVCF) across all samples for each reference and compared the impacts of five SV merging tools: BCFtools; Truvari; Jasmine; Naive 50% reciprocal-overlap; SURVIVOR. These tools use a variety of methods for SV comparison (Supplementary Table 2).

We’ve established the relationship between decreasing matching stringency and decreasing SV count. Here, BCFtools is the exact matching method and serves as an upper-limit to which we compare the other tools since it retains all redundant variants and therefore holds the maximum possible number of SVs. BCFtools produces 347,158 SVs for GRCh38 (80,322 DEL; 266,836 INS) and 329,937 SVs for chm13 (121,038 DEL; 208,899 INS). The lower-limit average allele frequency (AF) from BCFtools is 0.05. Using Truvari *anno repmask* and *anno numneigh*, we observe the highest number of SVs per-locus - and thus most likely in need of merging - are annotated as low complexity (average 8.5 SVs/locus) and simple repeats (6.7) (Supplementary Figure 4).

Relative to BCFtools, the merges have an average reduction in SV count of: Truvari 41%; Jasmine 59.8%; Naive 65.2%; SURVIVOR 77%. The largest difference in SV count reduction is between Truvari, which produces an average of 199,751 SVs, and SURVIVOR with 77,761. Broken down by SV type, this is a difference of ∼38.5K DEL and ∼83.5K INS. Additionally, the average AF observed in pVCFs is: Truvari 0.08; Jasmine 0.12; Naive 0.13; SURVIVOR 0.17. Therefore, choices in merging tools can cause an approximately 1.7x to 3.6x fold increase in AF. For details on SV count, average AF, and size distributions, see Supplementary Table 3, Supplementary Figure 5 & Supplementary Figure 6.

These patterns of SV count reduction and increased AF are not only present genome-wide, but also within genes. To highlight this, we used Truvari *anno bpovl* to identify SVs which intersect genes from Ensembl release-105[22] on GRCh38 (**Figure 3**). This shows that Truvari produces more variants at a lower average AF compared to the other tools which attempt to remove redundant alleles.

**Figure 3:**
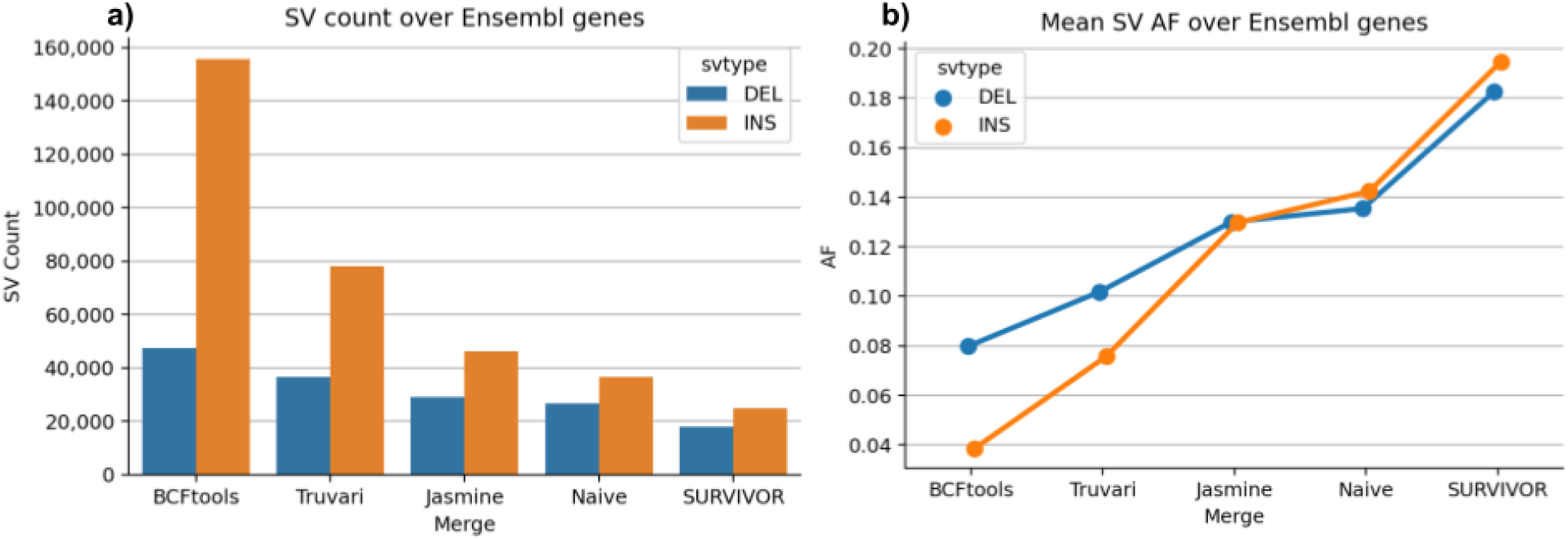
Merging strategies’ impact on pVCF number of SVs and their allele frequency over Ensembl genes a) Count of deletions and insertions produced by each merging strategy. b) Average allele frequency of SVs as merging leniency increases.

### Benchmarking pVCFs

GIAB recently published an expanded benchmark of challenging, medically relevant gene regions (CMRG)[23]. This includes 273 genes on GRCh38 which were resolved for structural variants in NA24385. In total, there are 216 SVs from NA24385 intersecting CMRG. Our SV calling pipeline identifies 2,363 SVs in CMRG regions across all individuals. Using Truvari *bench*, we assess how well merging tools are preserving variants by comparing non-reference-homozygous NA24385 sites in the pVCFs against CMRG. Because BCFtools only merges identical alleles and makes no attempt to remove redundancy, it has the highest possible recall with 201 TPs. However, of the tools that remove redundant variant representations, we again see that Truvari’s pVCF is best at preserving variants with one TP missing whereas the remaining tools over-merge and lose between 5 and 41 TPs (**Table 1**). Manual analysis found that Truvari’s single lost TP was inside the RNF213 gene at chr17:80274587 where two heterozygous insertions of length 538bp and 580bp with 96.2% sequence similarity were merged.

**Table 1:** Merges’ pVCF performance on GIAB CMRG SV benchmark of NA24385 for GRCh38. Hardy-Weinberg equilibrium (HWE) and excess heterozygosity (ExcHet) scores less than 0.05 were counted for true-positives across all 36 samples’ genotypes.

| Merge | TP | FP | FN | precision | recall | f1 | call cnt | gt_concordance | HWE<br>< 0.05 | ExcHet<br>< 0.05 |
| --- | --- | --- | --- | --- | --- | --- | --- | --- | --- | --- |
| BCFtools | 201 | 5 | 15 | 0.976 | 0.931 | 0.953 | 206 | 0.950 | 39 | 1 |
| Truvari | 200 | 5 | 16 | 0.976 | 0.926 | 0.950 | 205 | 0.950 | 17 | 4 |
| Jasmine | 196 | 10 | 20 | 0.951 | 0.907 | 0.929 | 206 | 0.949 | 17 | 4 |
| Naive | 171 | 22 | 45 | 0.886 | 0.792 | 0.836 | 193 | 0.947 | 26 | 18 |
| SURVIVOR | 160 | 28 | 56 | 0.851 | 0.741 | 0.792 | 188 | 0.956 | 24 | 20 |

Two metrics for evaluating the genotyping quality of variants are Hardy-Weinberg equilibrium (HWE) and excess heterozygosity (ExcHet) scores. Excluding variants with lower values of these scores is a common QC step in association studies[24]. Using NA24385 true-positives from each merge, we calculated the HWE and ExcHet across all 36 samples’ genotypes with the idea that fewer calls with lower scores (<0.05) indicates a higher quality merge (**Table 1**). The smallest proportions of TPs with low HWE are from Truvari and Jasmine results at 8.5% and 8.7%, respectively. The largest proportion is from the under-merged BCFtools results at 19.4% of all TPs. For ExcHet, we find up to 2% of BCFTools, Truvari, and Jasmine variants having low scores compared to 10.5% of Naive and 12.5% of SURVIVOR TPs.

### Assessing performance of merging tools

Beyond quantifying the differences of each merge, we need to assess how well they preserve measurably distinct alleles. The goal of SV merging is to identify redundant representations of alleles and consolidate their genotypes. Over-merging occurs when unique SV representations are falsely identified as being redundant. Ideally, a correct merge would retain all unique alleles while consolidating only truly redundant alleles.

One case where we expect an enrichment of redundant SV representations is in tandem repeat regions due to alignment ambiguities. Furthermore, we can classify all variants in a tandem repeat locus as representing unique or redundant expansions/contractions of the reference by running TandemRepeatFinder (**Figure 4a**, see methods). Since BCFtools performs exact matching and only identical alleles are consolidated, it preserves every input allele but fails to consolidate genotypes between redundant representations. Consequently, we can use BCFtools’ result as a baseline to which we compare in order to assess how many unique alleles are missing (over-merging) and how many redundant alleles remain (under-merging) in the SV merging tools’ results.

**Figure 4:**
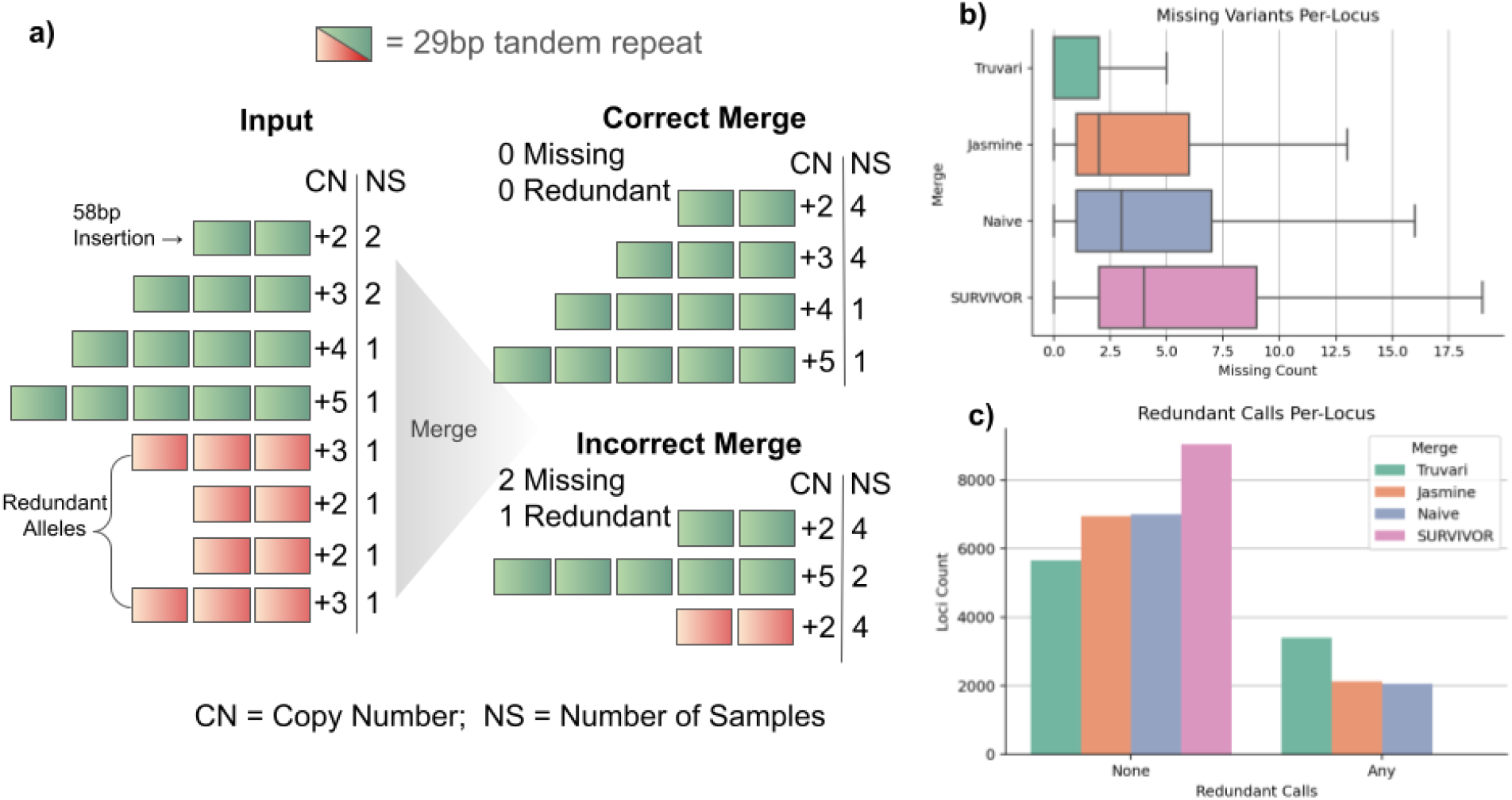
Investigation of tandem repeats to assess merging strategies’ performance a) illustration of a locus where eight insertion alleles (Input) have between +2 and +5 copies of a 29bp repeat across 10 samples. Four of the alleles are annotated as redundant representations (red) since they have a counterpart with an equal number of copies (green). A correct merge would preserve each of the unique alleles and remove all redundant alleles, leaving 0 missing and 0 redundant SVs in the locus. An incorrect merge removes two unique insertions (+3, +4) and leaves 1 redundant insertion. b) Boxplot of number of missing variants per-locus for each merging strategy. c) Barplot of the number of loci with None or Any redundant alleles post-merging.

We identified 20,207 tandem repeat loci with SVs. Of these tandem repeat loci, 9,056 (44%) have a different number of SVs reported from at least one merging tool. Thus, this subset of highly problematic regions were analyzed to assess the amount of missing alleles (**Figure 4b**) and redundant alleles (**Figure 4c**) in the merge tools’ pVCFs. Truvari had over-merging in 47.4% of loci (average 1.8 missing alleles per-locus). Jasmine and Naive had over-merging in 76.5% (4.4 alleles per-locus) and 79.6% (5.4) of loci, respectively. SURVIVOR, the most permissive SV merging tool, exhibited over-merging in 99.3% of loci, which averaged to 7.1 missing alleles per-locus. For loci with redundant alleles remaining Truvari produces 3,398 loci (37.5%) having at least 1 redundant allele compared to BCFtools, Jasmine 2,118 (23.3%), Naive 2,058 (22.7%), and SURVIVOR produces 13 loci (0.1%) with redundant alleles.

This analysis shows that using orthogonal information, we can objectively demonstrate that of the tools which attempt to identify and consolidate redundant allele representations, Truvari is performing best while other tools are over-merging more frequently, which in turn inflates AF and loses unique alleles.

## Discussion

In this paper we presented Truvari, a toolkit that enables merging, benchmarking, and annotations of SVs. We showed Truvari’s versatile applications to SV analysis and how it significantly improves the ability for researchers to accurately compare structural variants. We demonstrated this across an SV call-set for 36 haplotype-resolved long-read assemblies[16,17] by starting with the simplest case of SV merging and identifying identical alleles between haplotypes before progressively allowing more lenient matching with more permissive SV comparison thresholds. Naturally we observed the pattern of more lenient thresholds finding more matching SVs between haplotypes. As the problem of SV merging becomes more complex when merging between samples, we showed how Truvari’s approach outperforms other tools at preserving distinct alleles genome-wide, within genes, and in especially problematic tandem repeats. Throughout the project we measured performance of the SV calls with comparisons to GIAB SV benchmarks using Truvari[6,23].

While this research focused on SVs generated by long-read assemblies, Truvari is not restricted to input SVs produced from phased assemblies. Truvari’s flexibility allows it to be used on any VCF with SVs, even those generated by short reads. It’s important to note, however, that Truvari is currently most useful for ‘resolved’ SVs (i.e. DEL, INS, INV, and DUP).

Sequence resolved SVs are the desired future for the field with clear advantages to improve our understanding of the mechanisms and impacts of SVs. In this paper, we assessed the utility of sequence based comparisons and identified multiple locations where sequence similarity gave the final conclusion for if alleles should be considered equal. Nevertheless, Truvari currently uses parameterized thresholding to create decision boundaries for when SVs match.

Given the limited set of SVs that are fully resolved, it is unknown when alleles with high sequence similarity should remain unmatched. We assumed in this work that small sequence differences in alleles were due to sequencing errors, but some of these changes may represent biologically relevant differentiation. For example, in the tandem repeat performance assessment we identified Truvari as having the highest count of loci with redundant alleles remaining after merge. However, these ‘redundant’ alleles may be explained by point mutations in a copy of the tandem repeat such that TRF can still identify the repeat but the sequence is different enough to have biological consequences such as inhibiting the tandem repeat’s slipped-strand mispairing mechanisms[25]. If this is the case, these ‘redundant’ alleles shouldn’t be considered the same because the allele without accumulated point mutations may be more susceptible to further contraction/expansion of the tandem repeat than an allele with mutations. We are looking into these possibilities and investigating how dynamic thresholding can further improve Truvari’s SV comparison accuracy.

The overall importance of correct SV comparison is clear and we could showcase multiple improvements using Truvari. One of the most remarkable results is the shift in allele frequencies across the spectrum of merging tools. Figure 3 showed that other methods’ over-merging has a large impact on allele frequency, particularly for insertions. These differences have drastic implications on the interpretation of SVs across a population. This over-merging means that sequence differences between individuals are getting lost. Previous publications are indeed hinting towards a potential over-merge, but further investigations are needed as the number of studies investigating insertions, particularly those using long-reads, are small but rising[26–30]. Related to this is the overall number of SVs that one might assume in healthy human genomes (**Figure 3a**, Supplementary Figure 5). Using these phased assembly based SV call sets we could conclude that the number of SVs per human might be higher than expected which again highlights the importance of SVs and implies that the diversity of the human population may be underestimated.

## Conclusions

The choices made when performing SV comparison have important impacts on results. When SV comparison is too lenient, over-merging occurs, distinct alleles are lost, and metrics such as allele frequency are inflated. This research shows how Truvari’s method of leveraging multiple SV similarity metrics enables refined handling of SV comparison and a better approach to multi-sample SV analysis.

## Methods

### Truvari SV Comparison

Truvari’s core functionality (**Figure 1**) involves building a matrix of pairs of SVs and ordering the pairs to determine how each should be handled. To start, VCFs are consolidated using a ‘zipper’. This procedure opens sorted VCFs using pysam (a wrapper around htslib). The set of VCFs is then treated as a single stack where the ascending alphanumeric sorting of each chromosome and integer position is yielded by a generator. This zipped stack of variants is ‘chunked’ and all variants within a *chunksize* are grouped. The chunker is also responsible for variant filtering on properties such as size restrictions, reference location, or VCF FILTER as specified. Chunks are created between sets of variants where the maximum end position plus chunk size is greater than the start position of the next variant yielded from the zipper. The zipping and chunking infrastructure is reused for *bench* and *collapse*. Additional filtering parameters such as only comparing passing variants or those genotyped as being present (non-reference-homozygous) in a sample being analyzed are available and prevent calls from being used downstream.

The next step for the *bench* procedure is to build an NxM matrix of the base and comparison calls within a chunk of variants. If dimensions N or M are 0, all variants within the chunk are annotated as false-negatives (FN) or false-positives (FP). respectively. Each pair is then measured for similarity across multiple metrics to build a putative match.

Variants have the properties of start position (S), end position (E), length (L) and allele sequence (A). Deletion’s L(A) = E - S whereas Insertions have no span over the reference and length is simply L(A). Formal definitions of each metric follows:

#### Reference Distance

Variant’s positions are within the specified *refdist*

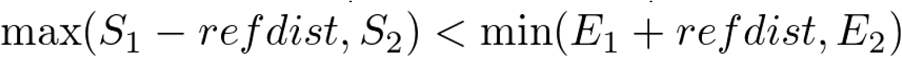

#### Reciprocal Overlap

Percent of overlapping bases over the maximum variant span.

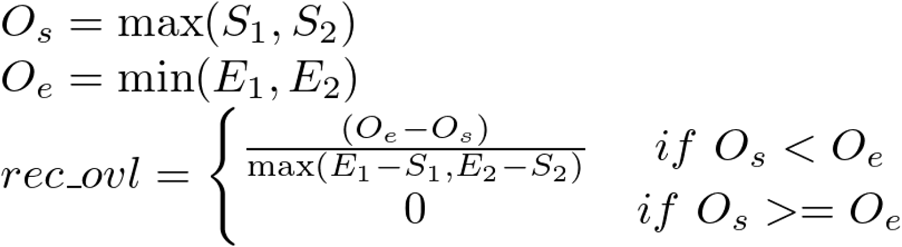

#### Size Similarity

Minimum variant length over the maximum variant length

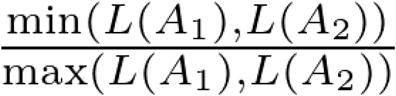

#### Sequence Similarity

Haplotype sequence similarity calculated with edlib[31].

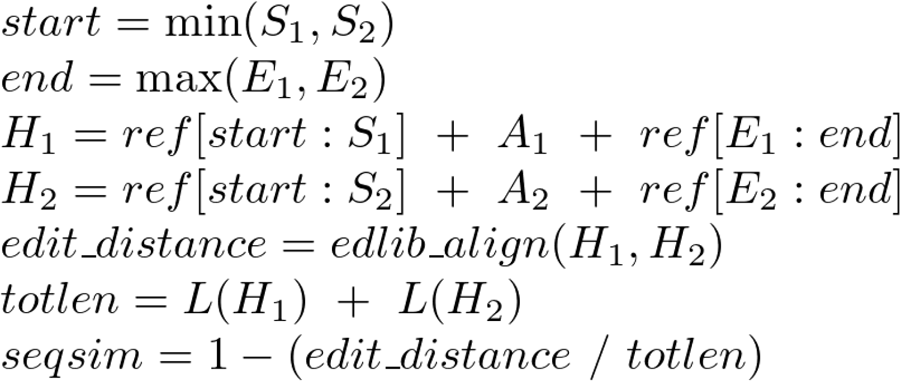

The reciprocal-overlap of sequence-resolved insertions, which have no physical span over the reference, is measured after the event’s boundaries are expanded by half the SV’s length upstream and downstream. To compute sequence similarity, the span of reference sequence between the two SV’s upstream-most start and downstream-most end is fetched and the sequence change of the SV is incorporated to create the shared sequence context of the calls. The two sequences are then aligned and similarity reported. The reciprocal-overlap, size similarity, and sequence similarity metrics are averaged to create a TruScore for ranking of putative matches. Each putative match is assumed to be valid until a comparison fails the thresholds/flags provided by the user. Additionally, the distance between the start and end breakpoints of the pair of calls is recorded for annotation purposes.

Once the matrix of putative matches is filled, it can be used to identify the best matches between the base and comparison calls. By default, only the single best match is searched for by raveling the 2D matrix into a 1D array and sorting the putative matches by their TruScore. Each match’s calls are checked to ensure they haven’t been used in a previous match. If neither have, the putative match with its state as determined by the thresholds is passed along to the output. If either call has been used previously, the match’s state is set to False and the unused base/comparison call in the pair is output as FP / FN respectively.

In some cases, a user may wish to allow variants to participate in more than one match. For example, one may expect multiple representations of an SV from a caller where there is only one inside the base variants. In this case, parsing the match matrix involves sorting each row and column independently by the TruScore such that the highest scoring match for each base and comparison call is reported.

For Truvari *collapse*, the same procedure to build matches is employed, however instead of a matrix of base/comparison, we have an NxN matrix of all calls within the chunk. Additionally, two more parameters are checked when building the match. If the user specified *--hap*, incompatible intra-sample genotypes are unable to be a valid match e.g. homozygous alternate calls in the same individual are not matched. The second parameter unique to Truvari *collapse* is *--chain*, which allows more flexibility around the *--refdist*. Chaining allows transitive matching such that two variants that don’t directly match but have a shared intermediate match are considered matching. After the matches have been built, each set of matching variants is sorted to determine which variant is kept in the output as the representative variant while the remaining are written to an extra VCF of collapsed variants. The options of which variant to keep from a set are: first - the most upstream variant; maxqual - the variant with the highest QUAL score; common - the variant with the highest minor allele count.

### Reference genomes

Human genome 19 (hg19), GRCh38, and telomere-to-telomere consortium chm13 v1.0 references were downloaded[18–20]. Alternate contigs were removed and variant calling was performed against only autosomes and the sex chromosomes X/Y.

### SV Calling

Previously published long-read, haplotype-resolved assemblies[16,17] were mapped with minimap2[32] version 2.17 and variants called with paftools, which is part of the minimap package. Minimap2 parameters used were “*-cx asm5 -t8 -k20 --secondary=no --cs ${ref} ${fasta}*” and paftools parameters “*-L10000*”. Three Individuals (HG00733, NA12878, NA24385) had assemblies created by both projects. In those cases, we chose to keep the assemblies generated by Garg et. al. as an attempt to increase heterogeneity of variants which would further test merging.

### Intra-sample haplotype merging

VCFs produced per-haplotype for each individual were merged using BCFtools v1.13[33]. A custom script consolidated genotypes to create a single SAMPLE column per-VCF. Truvari *collapse* v3.1 was run with *--hap* to prevent incompatible genotyped calls from being merged to produce the ‘strict’ intra-sample merge. Truvari *collapse* v3.1 parameters to produce the ‘loose’ merge “*--hap --pctsim 0*.*70 --pctsize 0*.*70 --refdist 1000*”. VCFs were converted to pandas DataFrames using Truvari *vcf2df* for analyses which can be recreated using the project’s github.

### RepeatMasker Classifications

Truvari *anno repmask* is a wrapper around RepeatMasker[34] that adds the annotation information into a VCF. For deletions, the REF sequence is run through RepeatMasker whereas for INS, the ALT sequence is used. For this study, a minimum RepeatMasker score of 250 was required to accept a reported annotation.

### Number of Neighbors

Truvari *anno numneigh* annotates entries in a VCF with how many other entries are within a specified distance as well as assigning an identifier for all variants within the same genomic region (i.e. neighborhood) as defined by the specified distance.

### GIAB Benchmarking

Comparisons to Genome in a Bottle consortium’s SVs v0.6 were performed against hg19[6] over the Tier1 regions. Comparisons to GIAB’s challenging, medically relevant genes (CMRG) SVs v1.0 were performed against GRCh38 over the resolved regions bed[35]. Truvari *bench* defaults were used. Hardy-Weinberg Equilibrium (HWE) and Excess Heterozygosity scores (ExcHet) were calculated using BCFtools +fill-tags.

### Inter-sample merging

Project-level VCFs were created using the per-sample VCFs generated by Truvari *collapse* with default parameters. BCFtools[33] version 1.13 had parameters “*-m none -0”*. Truvari *collapse* was run with *--chain* and default parameters. Jasmine[13] v1.14 was run with parameters “-*-output_genotypes --default_zero_genotype*”. SURVIVOR[11] v1.07 was run with parameters “*1000 1 1 0 1 50”*. Naive merging is performed by a custom script (available on the github) that merges variants with 500bp and with >= 50% reciprocal-overlap. Since insertion calls have no physical span over the reference (i.e. they exist between two reference bases), the Naive merging expands their boundaries to ±(SVLEN // 2). Allele frequencies within pVCFs were calculated using BCFtools *+fill-tags*. For most analyses, Truvari *vcf2df* was run to turn pVCFs into pandas DataFrame. Jupyter notebooks detailing steps of the analysis on github.

### Gene Intersection

Truvari *anno bpovl* was run to intersect pVCF entries to Ensembl release-v105[22] on GRCh38. This tool creates an Interval Tree for each range in the annotation file and checks variants’ intersection at the breakpoints as well as reporting if a variant is contained within or completely overlaps annotation file entries.

### Tandem Repeat Experiment

Truvari *anno trf* incorporates a wrapper around tandem-repeat finder (TRF)[36]. We ran Truvari *anno trf* to annotate all SVs on GRCh38 that intersected the SimpleRepeats track procured from UCSC Table Browser[29]. Each intersecting variant is used to alter the SimpleRepeat reference region to reconstruct the sample’s haplotype. TRF then detects the repeat sequence and copy number difference in an alternate allele relative to the reference (e.g. +5 copies of a 50bp repeat comprise a 250bp insertion). The longest tandem repeat found inside the altered sequence that’s shared with the reference annotations is reported as well as the copy-number difference of the variant compared to the reference track. Loci are identified using Truvari *anno numneigh* where variants within 1000bp are clustered. The BCFtools merge result serves as the baseline since it holds the maximum number of variants possible. Loci with at least one merging tool reporting a differing number of variants are analyzed. Each merging tools’ result is assessed per-locus and variant representations with unique copy-numbers missing or redundant representations remaining in the post-merge result are tallied.

## Abbreviations

SV: Structural Variant
QC: Quality Control
GIAB: Genome in a Bottle
VCF: Variant Call Format
pVCF: Project-level Variant Call Format
CMRG: Challenging Medically Relevant Genes
TRF: Tandem Repeat Finder

## Declarations

FJS received research funding from PacBio and Oxford Nanopore.

## Availability of data and materials

Truvari is available on https://github.com/ACEnglish/truvari Methods can be reproduced by code available on https://github.com/ACEnglish/truvaridata The README has details for commands to create all the data for analysis. The ‘manuscript/’ folder has post processing scripts and jupyter notebooks to recreate each figures and results in this paper. For full paths of all raw data used in this work, see the spreadsheet https://tinyurl.com/TruvariRawData This sheet also includes paths to the project’s final Truvari produced per-sample VCFs and pVCFs.

## Authors’ contributions

ACE performed software engineering of Truvari, data processing and analysis. ACE, FS did experimental design. ACE, VM, RG, GM, FS helped write the manuscript

## Acknowledgements

Adina Mangubat, Rob Flickenger, Niranjan Shekar, Nils McCarthy, Surabhi Maheshwari, Lisa Meed, and Jeremy Bruestle for help developing early versions of Truvari and generating data. Justin Zook, Nancy Hansen, and the Genome in a Bottle consortium for the initial inspiration for Truvari.

## Supplementary Figures/Tables

**Supplementary Figure 1.**
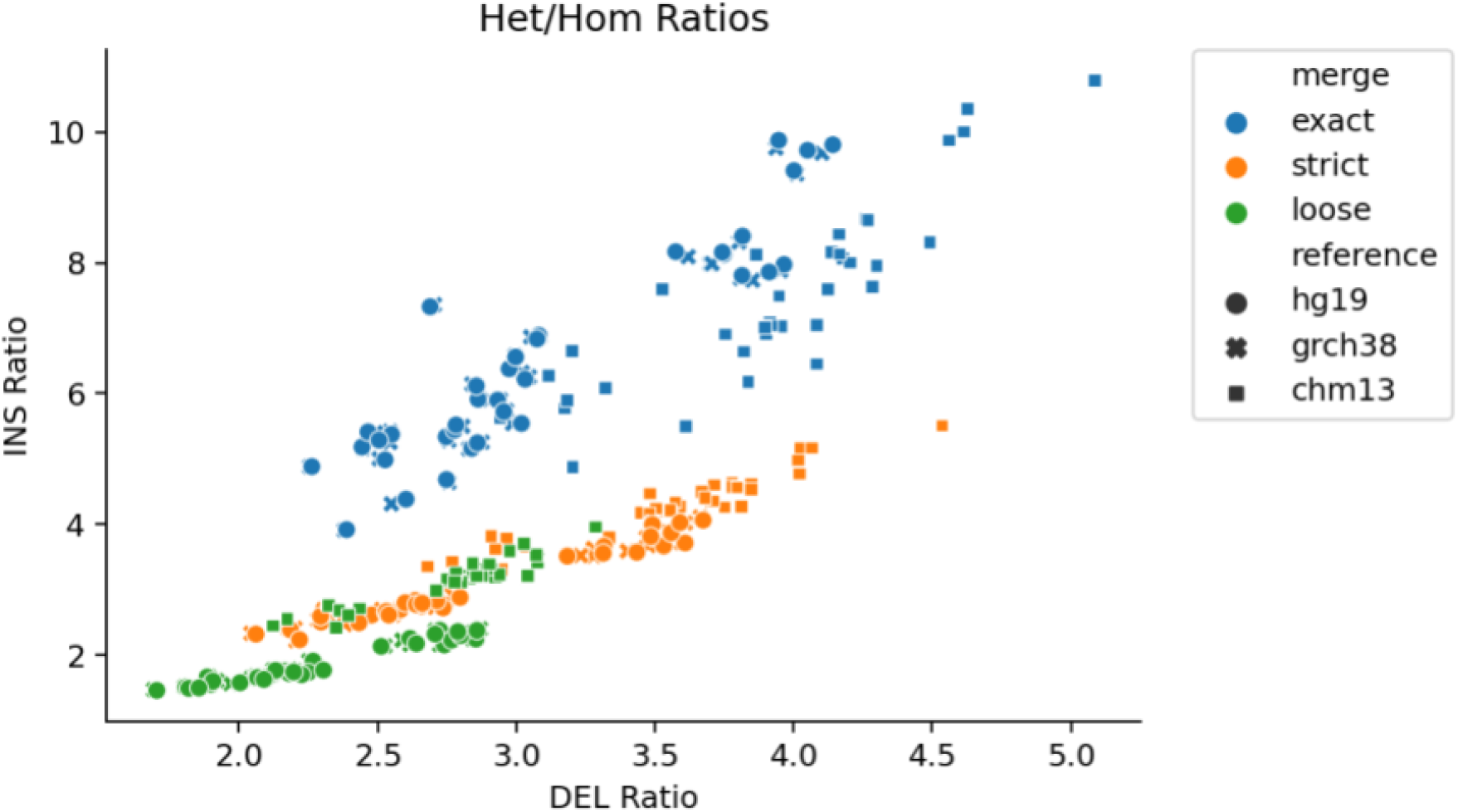
Het/Hom Ratios of Per-Sample VCFs by SVTYPE. Each point is a sample. Point colors are the intra-sample merge strategy. Shapes are references. As matching thresholds become more lenient, more heterozygous alleles find a counterpart and become homozygous, thus lowering the het/hom ratio. We see the ratios of INS (y-axis) dropping more quickly than DEL (x-axis).

**Supplementary Figure 2.**
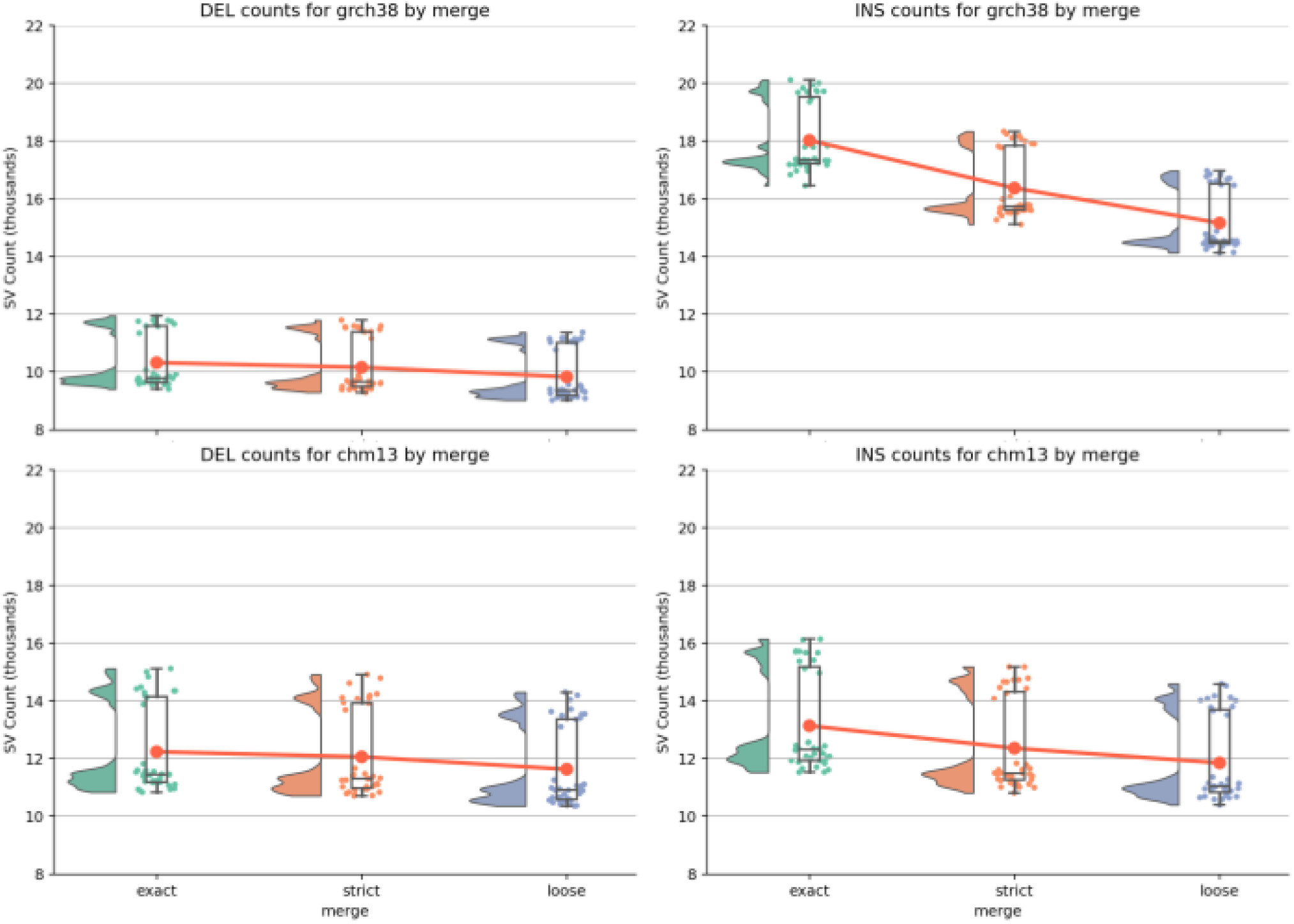
SV counts across inter-sample merges by SVTypes for GRCh38 and chm13. As matching thresholds become more lenient, more heterozygous alleles find a counterpart and become homozygous, thus lowering the SV count. We see a steeper decrease in INS counts than DELs, particularly for GRCh38.

**Supplementary Figure 3.**
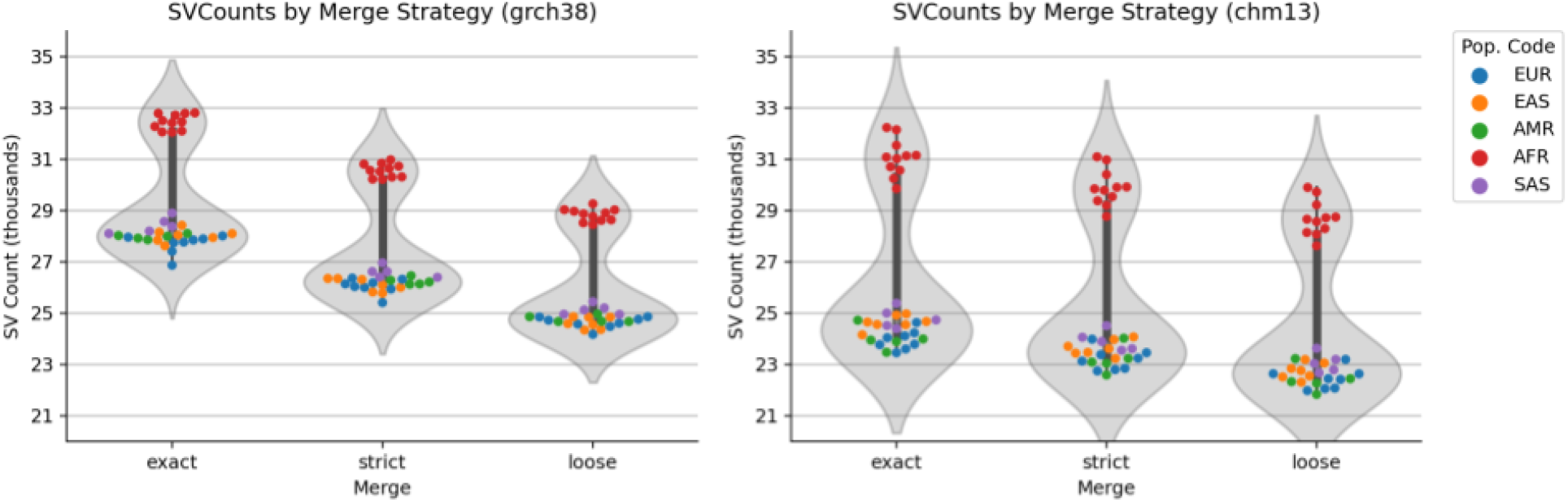
SV counts per-sample across merge strategies and references. Colors are sample’s population code. Samples from individuals of African ancestry have more SVs.

**Supplementary Figure 4.**
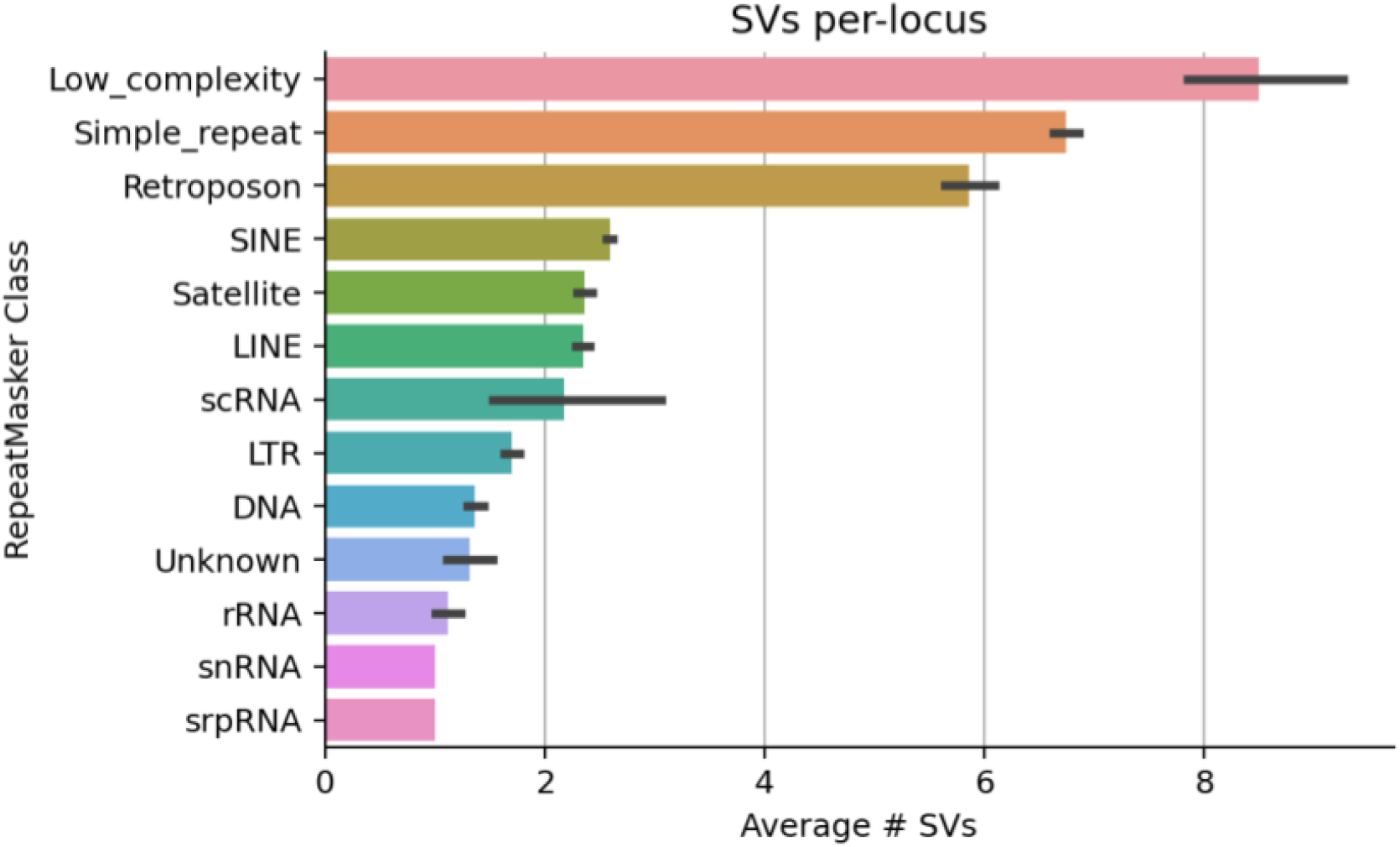
SVs per-locus by RepeatMasker class.

**Supplementary Figure 5.**
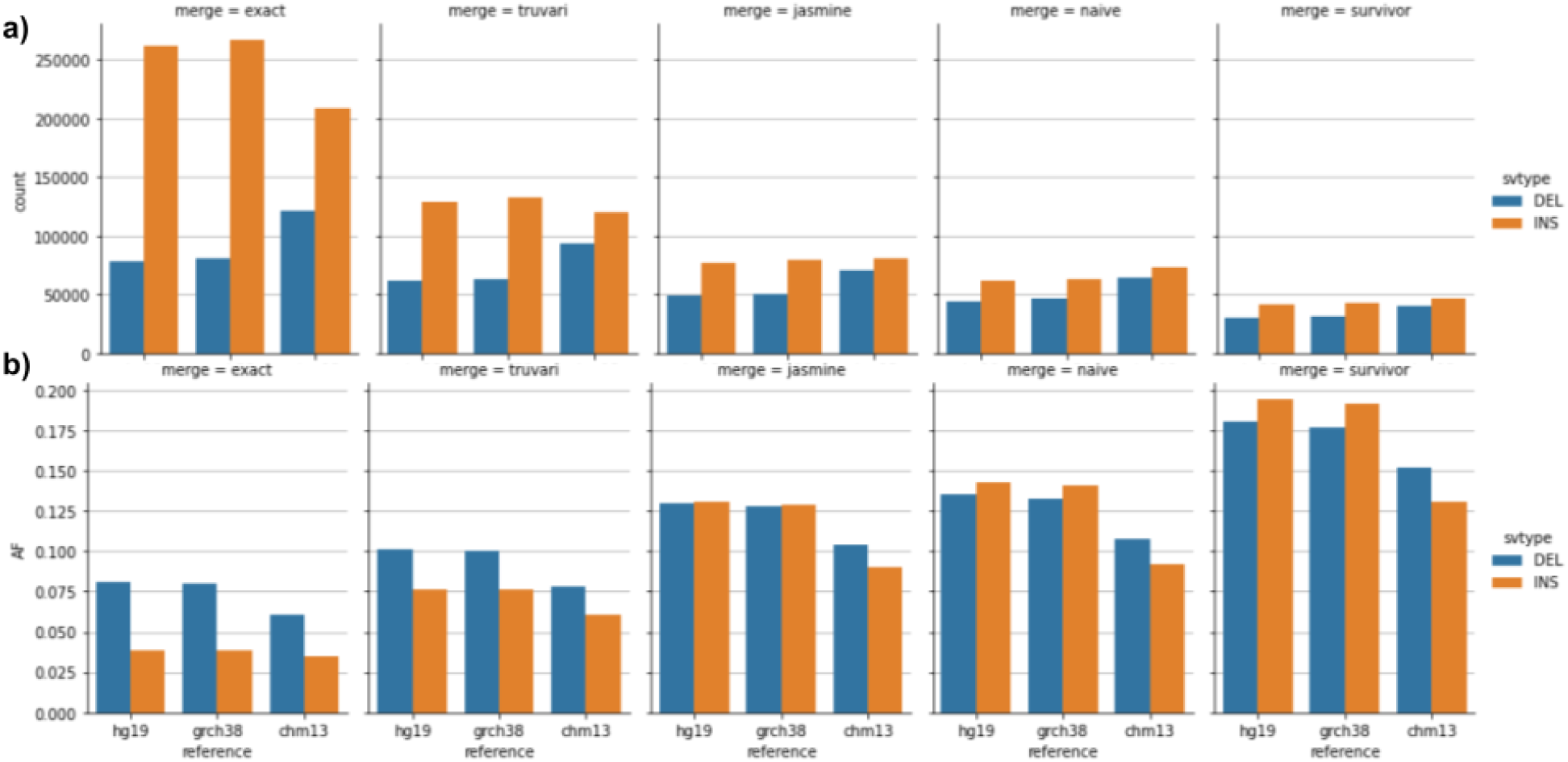
SVCount (a) and Allele-Frequency (b) for 5 SV merging tools (columns) across references (x-axis). We note very minor differences between hg19 and GRCh38.

**Supplementary Figure 6.**
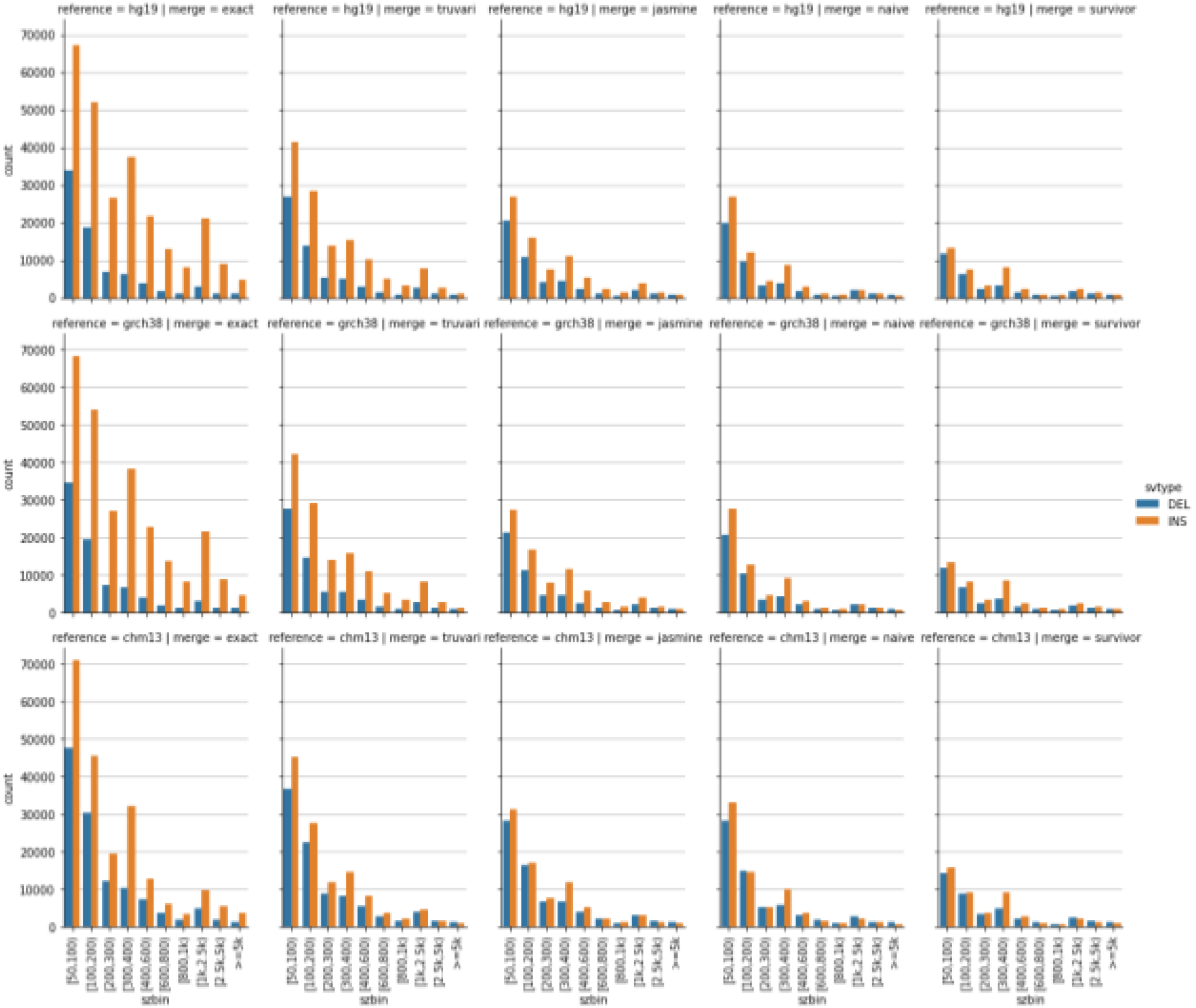
SVCount by size-bins (x-axis) for 5 SV merging tools (columns) across references (rows).

**Supplementary Table 1.** GIAB v0.6 Tier1 SVs performance of the three intra-sample merges on hg19

| Merge | TP-base | TP-call | FP | FN | precision | recall | f1 | base cnt | call cnt |
| --- | --- | --- | --- | --- | --- | --- | --- | --- | --- |
| exact | 9,036 | 10,185 | 933 | 605 | 0.916 | 0.937 | 0.927 | 9,641 | 11,118 |
| strict | 9,035 | 9,395 | 897 | 606 | 0.913 | 0.937 | 0.925 | 9,641 | 10,292 |
| loose | 9,028 | 9,058 | 834 | 613 | 0.916 | 0.936 | 0.926 | 9,641 | 9,892 |

**Supplementary Table 2.** Inter-Sample Merging tools’ versions and descriptions

| **Merge** | **Version** | **Estimated Stringency** | **Metrics** |
| --- | --- | --- | --- |
| BCFtools | 1.13 | Extreme | Identical position and sequence |
| Truvari | 3.1.0 | High | Type Matching, Reference Distance, Reciprocal Overlap, SizeSimilarity, SequenceSimilarity |
| Jasmine | 1.14 | Moderate | Type Matching, Position, Length |
| Naive | N/A | Moderate | Type Matching, Reciprocal Overlap |
| SURVIVOR | 1.07 | Low | Type Matching, Distance |

**Supplementary Table 3.**

| reference | merge | SV Count |  |  | Allele Frequency |  |  | Relative Count Reduction |  |  | AF Fold Increase |  |  | AF Relative to Truvari |
| --- | --- | --- | --- | --- | --- | --- | --- | --- | --- | --- | --- | --- | --- | --- |
|  |  | TOT | DEL | INS | TOT | DEL | INS | TOT | DEL | INS | TOT | DEL | INS |  |
| hg19 | exact | 339,870 | 78,036 | 261,834 | 0.05 | 0.08 | 0.04 | . | . | . | . | . | . | 0.6 |
| hg19 | truvari | 190,878 | 61,493 | 129,385 | 0.08 | 0.10 | 0.08 | 43.84% | 21.20% | 50.59% | 1.8 | 1.3 | 2.0 | . |
| hg19 | jasmine | 125,841 | 48,600 | 77,241 | 0.13 | 0.13 | 0.13 | 62.97% | 37.72% | 70.50% | 2.7 | 1.6 | 3.4 | 1.5 |
| hg19 | naive | 105,759 | 44,429 | 61,330 | 0.14 | 0.14 | 0.14 | 68.88% | 43.07% | 76.58% | 2.9 | 1.7 | 3.7 | 1.7 |
| hg19 | survivor | 71,845 | 30,372 | 41,473 | 0.19 | 0.18 | 0.19 | 78.86% | 61.08% | 84.16% | 3.9 | 2.2 | 5.1 | 2.2 |
| grch38 | exact | 347,158 | 80,322 | 266,836 | 0.05 | 0.08 | 0.04 | . | . | . | . | . | . | 0.6 |
| grch38 | truvari | 195,705 | 63,418 | 132,287 | 0.08 | 0.10 | 0.08 | 43.63% | 21.05% | 50.42% | 1.7 | 1.3 | 2.0 | . |
| grch38 | jasmine | 129,744 | 50,278 | 79,466 | 0.13 | 0.13 | 0.13 | 62.63% | 37.40% | 70.22% | 2.7 | 1.6 | 3.4 | 1.5 |
| grch38 | naive | 109,379 | 46,087 | 63,292 | 0.14 | 0.13 | 0.14 | 68.49% | 42.62% | 76.28% | 2.9 | 1.7 | 3.7 | 1.6 |
| grch38 | survivor | 74,288 | 31,560 | 42,728 | 0.19 | 0.18 | 0.19 | 78.60% | 60.71% | 83.99% | 3.9 | 2.2 | 5.0 | 2.2 |
| chm13 | exact | 329,937 | 121,038 | 208,899 | 0.04 | 0.06 | 0.04 | . | . | . | . | . | . | 0.7 |
| chm13 | truvari | 212,670 | 92,996 | 119,674 | 0.07 | 0.08 | 0.06 | 35.54% | 23.17% | 42.71% | 1.5 | 1.3 | 1.7 | . |
| chm13 | jasmine | 151,975 | 70,696 | 81,279 | 0.10 | 0.10 | 0.09 | 53.94% | 41.59% | 61.09% | 2.2 | 1.7 | 2.6 | 1.4 |
| chm13 | naive | 137,594 | 64,361 | 73,233 | 0.10 | 0.11 | 0.09 | 58.30% | 46.83% | 64.94% | 2.2 | 1.8 | 2.6 | 1.5 |
| chm13 | survivor | 87,150 | 40,554 | 46,596 | 0.14 | 0.15 | 0.13 | 73.59% | 66.49% | 77.69% | 3.2 | 2.5 | 3.7 | 2.1 |
| Average | exact | 338,988 | 93,132 | 245,856 | 0.05 | 0.07 | 0.04 | . | . | . | . | . | . | 0.6 |
| Average | truvari | 199,751 | 72,636 | 127,115 | 0.08 | 0.09 | 0.07 | 41.00% | 21.80% | 47.91% | 1.7 | 1.3 | 1.9 | . |
| Average | jasmine | 135,853 | 56,525 | 79,329 | 0.12 | 0.12 | 0.12 | 59.85% | 38.91% | 67.27% | 2.5 | 1.6 | 3.1 | 1.5 |
| Average | naive | 117,577 | 51,626 | 65,952 | 0.13 | 0.13 | 0.13 | 65.22% | 44.17% | 72.60% | 2.7 | 1.7 | 3.3 | 1.6 |
| Average | survivor | 77,761 | 34,162 | 43,599 | 0.17 | 0.17 | 0.17 | 77.02% | 62.76% | 81.95% | 3.6 | 2.3 | 4.6 | 2.2 |

